# Kar4 is Required for the Normal Pattern of Meiotic Gene Expression

**DOI:** 10.1101/2023.01.29.526097

**Authors:** Zachory M. Park, Matthew Remillard, Mark D. Rose

## Abstract

Kar4p, the yeast homolog of the mammalian methyltransferase subunit METTL14, is required for the initiation of meiosis and has at least two distinct functions in regulating the meiotic program. Cells lacking Kar4p can be driven to sporulate by co-overexpressing the master meiotic transcription factor, *IME1*, and the translational regulator, *RIM4*, suggesting that Kar4p functions at both the transcriptional and translational level to regulate meiosis. Using microarray analysis and RNA sequencing, we found that *kar4*Δ/Δ mutants have a largely wild type transcriptional profile with the exception of two groups of genes that show delayed and reduced expression: (1) a set of Ime1p-dependent early genes as well as *IME1*, and (2) a set of late genes dependent on the mid-meiotic transcription factor, Ndt80p. The early gene expression defect is rescued by overexpressing *IME1*, but the late defect is only suppressed by overexpression of both *IME1* and *RIM4*. Mass spectrometry analysis identified several genes involved in meiotic recombination with strongly reduced protein levels, but with little to no reduction in transcript levels in *kar4*Δ/Δ after *IME1* overexpression. The low levels of these proteins were rescued by overexpression of *RIM4* and *IME1*, but not by the overexpression of *IME1* alone. These data expand our understanding of the role of Kar4p in regulating meiosis and provide key insights into a potential mechanism of Kar4p’s later meiotic function that is independent of mRNA methylation.

**Author Summary:** Kar4p is required at two stages during meiosis. Cells lacking Kar4p have a severe loss of mRNA methylation and arrest early in the meiotic program, failing to undergo either pre-meiotic DNA synthesis or meiotic recombination. The early block is rescued by overexpression of the meiotic transcription factor, *IME1*. The *kar4*Δ/Δ cells show delayed and reduced expression of a set of Ime1p-dependent genes expressed early in meiosis as well as a set of later genes that are largely Ndt80p-dependent. Overexpression of *IME1* rescues the expression defect of these early genes and expedites the meiotic program in the wild type S288C strain background. However, *IME1* overexpression is not sufficient to facilitate sporulation in *kar4*Δ/Δ. Completion of meiosis and sporulation requires the additional overexpression of a translational regulator, *RIM4*.

Analysis of *kar4*Δ/Δ’s proteome during meiosis with *IME1* overexpression revealed that proteins important for meiotic recombination have reduced levels that cannot be explained by equivalent reductions in transcript abundance. *IME1* overexpression by itself rescues the defect associated with a catalytic mutant of Ime4p, implying that the early defect reflects mRNA methylation. The residual defects in protein levels likely reflect the loss of a non-catalytic function of Kar4p, and the methylation complex, which requires overexpression of *RIM4* to suppress.

## Introduction

Meiosis is a highly conserved eukaryotic cell differentiation process that begins with a single diploid cell and culminates in the production of four haploid cells. In the budding yeast, *Saccharomyces cerevisiae*, meiosis occurs in response to environmental stress and results in the production of four haploid spores (gametes) contained in a single ascus.

The yeast meiotic program is initiated when diploid cells are exposed to conditions lacking nitrogen and a fermentable carbon source. Nutrient signaling is coupled with ploidy sensing mediated by *MAT*a1/α2 leading to expression of the early meiotic transcription factor, Ime1p (Neiman 2011). Ime1p initiates the expression of a set of genes required for the initiation and early steps of the meiotic program, including pre-meiotic S-phase and meiotic recombination. Within Ime1p’s regulon is the meiotic protein kinase Ime2p. Ime2p functions to initiate pre-meiotic S-phase, activate the middle meiotic transcription factor, Ndt80p, and turn off *IME1* expression, as well as other regulatory roles that are essential for the proper completion of meiosis (Kassir, Granot et al. 1988, Dirick, Goetsch et al. 1998, Clifford, Marinco et al. 2004, Sedgwick, Rawluk et al. 2006, Holt, Hutti et al. 2007, Ahmed, Bungard et al. 2009, Corbi, Sunder et al. 2014). Activation of Ndt80p induces the expression of genes required for completion of the meiotic divisions and spore maturation (Pak and Segall 2002, Ahmed, Bungard et al. 2009, Shin, Skokotas et al. 2010, Neiman 2011, van Werven and Amon 2011, Winter 2012). In addition to these three key meiotic regulators, the RNA-binding translational regulator Rim4p has also emerged as an important regulator of the meiotic program. Rim4p has been shown to have two functions: 1) it activates the expression of early meiotic genes, including *IME2* (Soushko and Mitchell 2000, Deng and Saunders 2001), and 2) it blocks the translation of mid-late meiotic genes including *CLB3* that are transcribed before their protein products are required (Berchowitz, Gajadhar et al. 2013). The block to translation is mediated by the sequestration of the regulated mRNAs into amyloid-like aggregates formed through Rim4p’s C-terminal low-complexity domain (Berchowitz, Kabachinski et al. 2015). The aggregates are dissolved by Ime2p phosphorylation, which releases the bound mRNAs making them accessible to the translation machinery. Ime2p phosphorylation also targets Rim4p for degradation via autophagy (Jin, Zhang et al. 2015, Carpenter, Bell et al. 2018, Wang, Zhang et al. 2020).

Another key mechanism of regulation during meiosis is mRNA m^6^A methylation. Among the most abundant mRNA modifications, m^6^A methylation is widespread among eukaryotes.Methylation is catalyzed by a trimeric complex, composed of METTL3, METTL14, and WTAP, which is highly conserved across eukaryotes (Bujnicki, Feder et al. 2002, Wang, Doxtader et al. 2016, Wang, Feng et al. 2016). In yeast, the complex was initially identified as containing Ime4p (the ortholog of METTL3), Mum2p (the ortholog of WTAP), and Slz1p (Clancy, Shambaugh et al. 2002, Agarwala, Blitzblau et al. 2012). However, work described in concurrent papers (Park, Sporer et al. 2023 (companion manuscript) and Ensinck, Maman et al. 2023 manuscript in preparation) showed that Kar4p, the ortholog of METTL14, is also part of the catalytic complex and required for mRNA methylation. In yeast, mRNA methylation levels peak early in meiosis before the induced expression of *NDT80* and initiation of the first meiotic division (Agarwala, Blitzblau et al. 2012). Methylation is present mainly around 3’ UTRs and the methylated transcripts are enriched on translating ribosomes. The methylated transcripts are enriched from genes involved in early stages of meiosis including DNA replication and recombination. Of particular importance, the transcripts of key regulators of meiosis have been reported to be methylated including *IME1, IME2*, and *RME1* (Bodi, Button et al. 2010, Schwartz, Agarwala et al. 2013, Bodi, Bottley et al. 2015). *RME1* encodes the main transcriptional repressor of *IME1*; methylation of *RME1* transcripts is associated with more rapid turnover. The reduction in *RME1* transcript levels leads to reduced Rme1 protein production and increased expression of *IME1*; increased *IME1* subsequently licenses cells to initiate the meiotic program. Remarkably, Ime4p has also been implicated in meiotic functions that are independent of mRNA methylation (Bushkin, Pincus et al. 2019). Taken together, transcriptional, post-transcriptional, and post-translational mechanisms of regulation are all required to ensure proper completion of the meiotic program.

In our concurrent paper, we show that the yeast karyogamy protein Kar4p is required early in meiosis before pre-meiotic S-phase and has two distinct functions in meiosis, termed Mei and Spo. These two functions are distinct from its function during yeast mating (the Mat function), where it acts with the key mating transcription factor, Ste12p, to promote the transcription of genes required for late steps during mating (Kurihara, Beh et al. 1994, Kurihara, Stewart et al. 1996, Lahav, Gammie et al. 2007). Kar4p is required for efficient mRNA methylation during meiosis, resulting in higher Rme1p levels in *kar4*Δ/Δ and lower Ime1p levels, causing an early defect in meiosis. Consistent with this, hypomorphic alleles of *RME1* permit pre-meiotic S-phase in *kar4*Δ/Δ, although not spore formation. Ectopic overexpression of *IME1* also suppresses the early Mei-*kar4*Δ/Δ defect. However, the additional overexpression of *RIM4* is required to suppress the Spo^-^ defect and allow sporulation in *kar4*Δ/Δ. The Mei^-^ defect is associated with Kar4p’s role in mRNA methylation given that methylation acts upstream of *IME1* and is similar to that of an *IME4* catalytic mutant. That Rim4p overexpression is required to suppress the Spo^-^ defect, but is not needed to suppress the *IME4* catalytic mutation, suggests that Kar4p, like Ime4p, may also regulate meiosis via a mechanism that is independent of mRNA methylation.

Here, we use transcriptomic and proteomic analysis to examine how Kar4p impacts the meiotic transcriptional landscape and to determine the nature of the Spo-meiotic defect.Microarray and RNA-seq data show that *kar4*Δ/Δ mutants have both an early transcriptional defect, which is rescued by overexpressing *IME1*, and a late transcriptional defect, which is rescued by additionally overexpressing *RIM4*. The late transcriptional defect largely involves genes within Ndt80p’s regulon, and we see a strong defect in *NDT80* expression in *kar4*Δ/Δ. In addition, mass spectrometry (MS) identified a subset of proteins that are disproportionately reduced relative to their transcript levels in *kar4*Δ/Δ, even with overexpressed *IME1*. The protein levels were restored by the additional overexpression of *RIM4*, without correlated changes in the level of gene expression. This suggests that *RIM4* overexpression likely impacts the efficiency of translation of these transcripts as opposed to increasing their overall levels. Taken together, these findings support a model in which Kar4p acts early through regulating *IME1* expression and has a later function that appears to be at least partially upstream of Ndt80p, which functions to positively regulate the translation of a set of transcripts required during various stages of meiosis.

## Results

### Meiotic Transcriptional Profile of *kar4*Δ/Δ

The requirement for Kar4p in mRNA methylation during meiosis, that mRNA methylation acts upstream of *IME1*, and that *IME1* overexpression bypasses the Mei-*kar4*Δ/Δ defect all suggest that there should be differences in the meiotic transcriptional profiles between wild type and *kar4*Δ/Δ cells. To determine if there is a transcriptional defect in *kar4*Δ/Δ cells during meiosis, we used microarrays to measure mRNA abundance in wild type and *kar4*Δ/Δ strains over the first 16 hours of meiosis. RNA sequencing was also conducted across several time points to validate the microarray data. The data show remarkably similar expression profiles between wild type and *kar4*Δ/Δ. However, starting at 7 hours there are 3 prominent gene clusters with lower expression in *kar4*Δ*/*Δ. The genes in these clusters are implicated in mid- and late-meiotic function, including cell cycle regulation, spindle assembly, and sporulation, based on gene ontology (GO) term analysis (Fig 1A-B, S Fig 1). Although these clusters are noteworthy, it is likely that they are indirect effects. First, the initial *kar4*Δ*/*Δ defect occurs before premeiotic S-phase, with DNA replication, meiotic recombination, and sporulation absent in *kar4*Δ*/*Δ cells. Second, the defect of Mei^-^ Kar4p mutants can be suppressed by over-expressing *IME1*, which acts at the initiation of meiosis, long before expression of the mid- and late-meiosis genes. Thus, it is likely that the large changes in late gene expression reflect the consequences of an earlier, more subtle defect.

**Fig 1.**
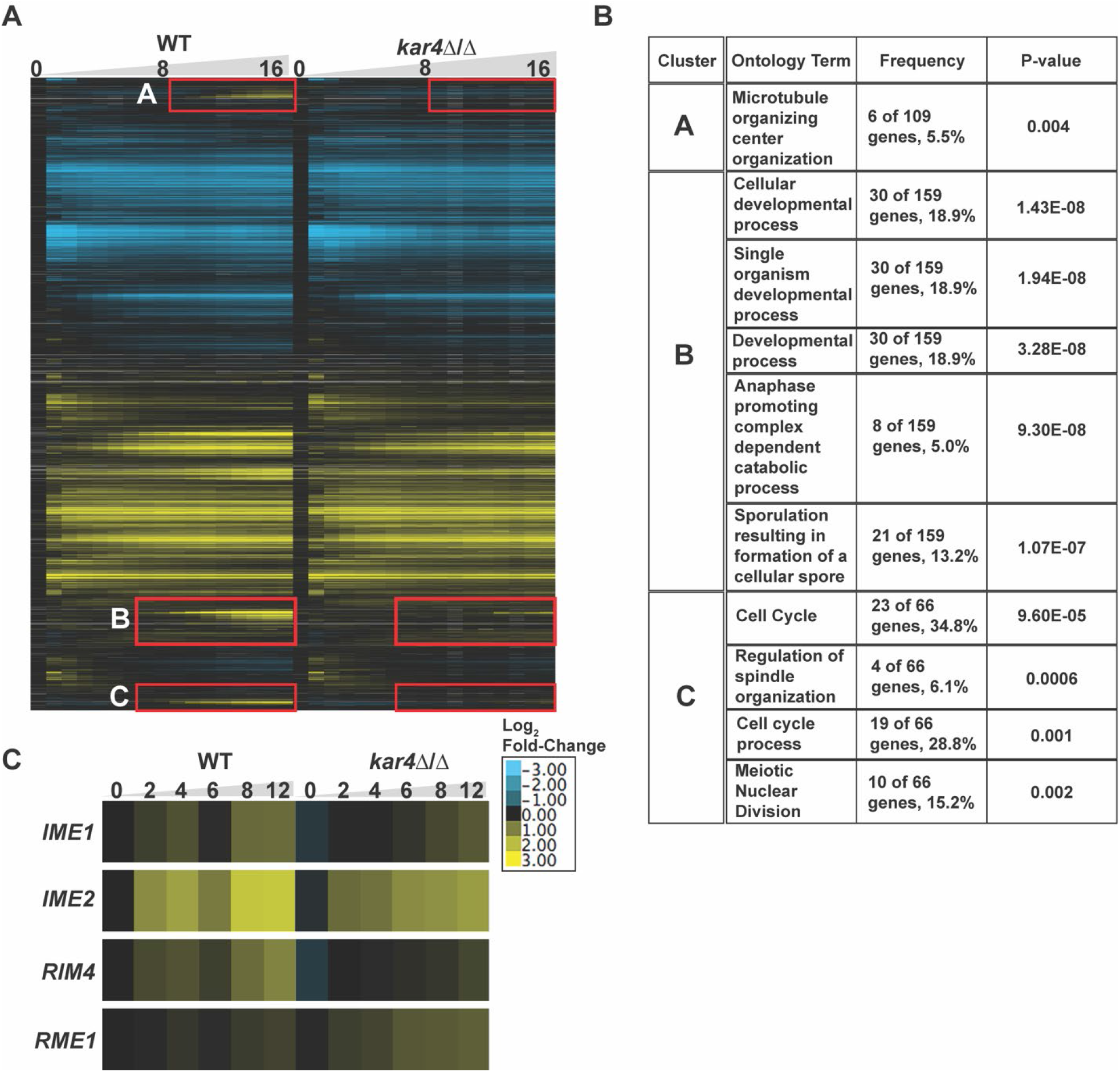
The meiotic transcriptome of *kar4*Δ/Δ. (A) Heatmap of microarray data across the first 16 hours post induction of meiosis in wild type and *kar4*Δ/Δ. Expression was normalized to wild type pre-induction of sporulation (t=0). Genes were clustered in Cluster3.0 and the heatmaps were constructed with Java TreeView. Red boxes highlight three clusters of altered late gene expression. (B) Table of the top gene ontology terms of the three clusters (A, B, and C) of impacted genes in *kar4*Δ/Δ. (C) Heatmap of RNA-seq data showing the expression level of key meiotic regulators in wild type and *kar4*Δ/Δ.

RNA-seq revealed that *IME1* transcript levels as well as the transcript levels of other key early meiotic regulators, *RIM4* and *IME2*, showed delayed and reduced expression in *kar4*Δ/Δ. In contrast, there was increased expression of the negative transcriptional regulator of *IME1, RME1*, across the time course (Fig 1C). Pervious work demonstrated that mRNA methylation destabilizes *RME1* transcripts and leads to increased expression of *IME1* (Bushkin, Pincus et al. 2019). Consistent with this, Ime1p levels are lower and Rme1p levels are higher in *kar4*Δ/Δ during meiosis and reducing Rme1p levels permits pre-meiotic S-phase in *kar4*Δ/Δ (see companion manuscript). Thus, the reduction in *IME1* expression may be due to the loss of negative regulation on *RME1* transcripts in *kar4*Δ/Δ. However, it is surprising that a relatively small effect on *IME1* levels in *kar4*Δ/Δ totally blocks cells from meiotic entry. One possibility is that *kar4*Δ/Δ has a stronger impact on a small fraction of Ime1p-dependent genes that are required for early stages of meiosis. To address this issue, we sought to identify a more specific set of Ime1p-dependent meiotic genes and examine their expression in *kar4*Δ/Δ.

### *KAR4* is Required for a Subset of *IME1*-Dependent Genes

The direct transcriptional targets of Ime1p are not well characterized. [YEASTRACT](http://www.yeastract.com/), a curated repository of yeast transcription factors and target genes, contains over 122 “experimentally” identified targets and 1071 putative targets based on the upstream activation sequence (UAS) “TTTTCHHCG” (Bowdish and Mitchell 1993, Monteiro, Oliveira et al. 2020). Many of the experimentally defined targets have been classified only on whether their expression is Ime1p-dependent, meaning some could be indirect. With nearly 20 percent of the yeast genome listed as possible genes of interest, this dataset was not useful for identifying Ime1p-dependent genes potentially impacted by Kar4p. in our transcriptome data. We therefore sought to create a more specific set of *IME1*-dependent genes using a strain with *IME1* under the control of the estradiol-inducible P_Z3EV_ (referred to as Pzev) promoter to overexpress *IME1* and identify genes that are rapidly induced under sporulation conditions (McIsaac, Gibney et al. 2014). Specifically, we compared gene expression profiles of Pzev-*IME1* strains after 2 hours in sporulation media with and without estradiol. Our data show *IME1* expression in the estradiol sample, with no *IME1* expression in the control sample.

We identified 236 genes with greater than 2-fold increased expression after *IME1* induction. However, it is possible that strong induction of *IME1* caused expression of off-target genes. Therefore, we compared the Pzev-*IME1* induction data with the wild type *pIME1* meiotic transcription profile to identify the subset of Pzev-*IME1* induced genes that show increased expression during normal meiosis. This criterion resulted in a list of 140 *IME1*-induced genes.

The set of 140 *IME1*-induced genes contains only 13 of the 126 identified targets and 53 of the 1071 putative targets in YEASTRACT. One possible difference is that the YEASTRACT gene list does not consider the timing of transcription or indirect targets over the entire meiotic program, whereas our analysis resulted in a list of initial Ime1p targets after 2 hours of induction.

To assess the quality of our Ime1p-dependent gene list, we performed GO-term analysis on our list and the YEASTRACT *IME1-*regulated gene list. Analysis of our 140 *IME1*-induced genes returned 65 GO terms; “meiotic nuclear division” and “meiotic cell cycle” are the top two results (P-value 1.1 × 10^−30^ and 2.2 × 10^−29^ respectively) (Fig 2A). In contrast, analysis of the YEASTRACT *IME1* regulated genes resulted in 17 GO-terms, suggesting poor specificity and less robust enrichment. The top result was “DNA Recombination” (P-value 8.2 10^−10^) and the GO term “meiotic cell cycle” is the seventh result (P-value 0.001). Our gene list captured more than three times the number of significant GO terms with increased specificity and coverage. The results of our GO term analysis support these data as a more defined set of genes that are directly regulated by Ime1p.

**Fig 2.**
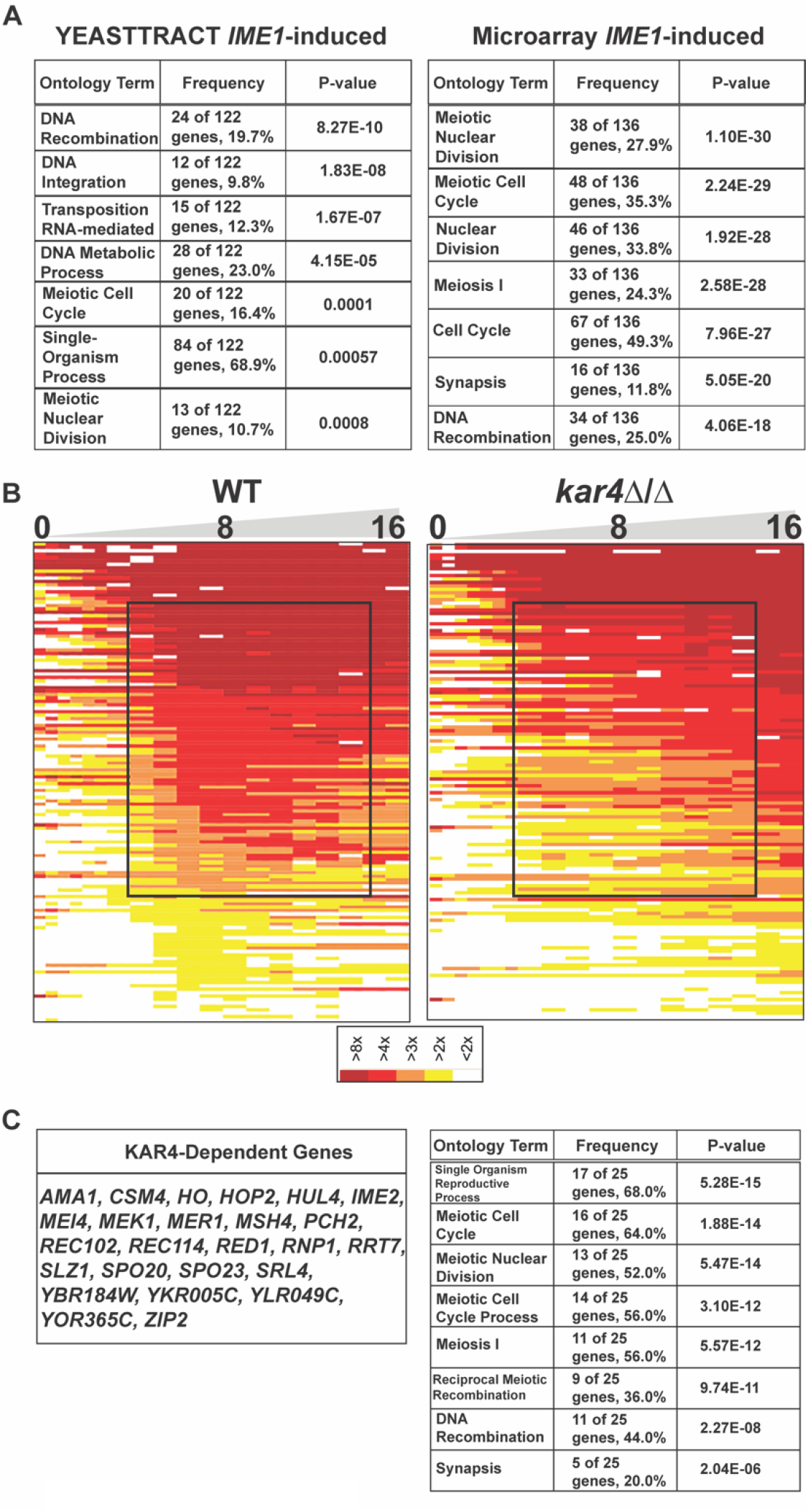
Kar4p is required for the expression of Ime1p dependent genes. (A) Gene Ontology results of the YEASTTRACT list of Ime1p dependent genes (Left) and of the gold-standard list of Ime1p dependent genes found in this study (Right). (B) Heatmap of microarray data showing the expression profile of Ime1p dependent genes in wild type and *kar4*Δ/Δ. The black box highlights the reduced and delayed expression of a set of these genes in *kar4*Δ/Δ. (C) List of Kar4p dependent genes that are also Ime1p dependent (Left) and the top gene ontology terms of those genes (Right).

Using this list, we examined whether *kar4*Δ/Δ altered the expression of the Ime1p-induced genes. To visualize the expression dynamics of these Ime1p-dependent genes, we ordered them by their average expression between 6 and 13 hours and found that in wild type cells there were two waves of *IME1*-dependent gene expression. In the first wave, between 0 and 3.5 hours, 9 genes were strongly expressed. The second wave occurred between 6 and 13 hours and 38 genes were strongly expressed. The early expressing group of genes showed no differential expression between wild type and *kar4*Δ/Δ. However, a large group of genes expressed between 6 and 13 hours appeared to show severely delayed and reduced expression in *kar4*Δ/Δ cells compared to wild type (Fig 2B). We defined *KAR4*-dependence as genes whose average expression from 1 to 3.5 (early I) or 6 to 13 hours (early II) is 2-fold or greater in wild type. Using those criteria, we identified 25 genes with a 2-fold or greater defect in expression in *kar4*Δ/Δ compared to wild type, although there were many more that showed smaller effects (Fig 2C).

Previous work has shown that there is an early burst of *IME1* expression that is independent of mRNA methylation, but the sustained increase in *IME1* expression that occurs as the cells continue through meiosis requires mRNA methylation (Bushkin, Pincus et al. 2019).

The lack of an effect on the expression of the early set of Ime1p-dependent genes in *kar4*Δ/Δ is consistent with this early burst of methylation-independent *IME1* expression driving the expression of those genes. The delayed and reduced expression of the later set of genes in *kar4*Δ/Δ is consistent with the requirement for mRNA methylation in the normal expression of *IME1* and genes in its regulon as the cells continue to move through meiosis. That these genes are eventually expressed in *kar4*Δ/Δ may be because mRNA methylation is not totally lost and/or that enough Ime1p is eventually made to drive their expression. Taken together, these data support a role for Kar4p in regulating the progression of cells into meiosis via regulation of *IME1* and genes in its regulon, consistent with findings in our companion manuscript showing lower levels of *IME1* transcript and protein in *kar4*∆/∆.

### *IME1* Overexpression Suppresses the *kar4*Δ/Δ Early Transcript Abundance Defect

*IME1* overexpression partially rescued the *kar4*Δ/Δ meiotic defect allowing pre-meiotic S-phase and meiotic recombination (see companion manuscript). To determine whether *IME1* overexpression rescues the reduction in Ime1p-dependent gene expression we examined gene expression in wild type and *kar4*Δ/Δ strains containing Pzev-*IME1*. As expected, estradiol induction resulted in rapid induction of the *IME1*-induced genes (Fig 3A), earlier than in cells with the wild type *IME1* promoter. In *kar4*Δ/Δ, overexpression of Ime1p was sufficient to rescue the early transcript abundance defect of the Ime1p-dependent genes (Fig 3A), supporting the hypothesis that *IME1* overexpression bypasses the requirement for Kar4p in establishing the early meiotic transcriptional profile. However, overexpression of Ime1p did not suppress the defect in gene expression observed for the three clusters of late genes (Fig 3B). Impacted genes are involved in later meiotic processes such as the meiotic divisions and spore formation and largely fall under the control of the mid-meiotic transcription factor Ndt80p. Consistent with a defect in Ndt80p-derependent regulation, the transcript abundance of *NDT80* in *kar4*Δ/Δ was reduced over 2-fold compared to wild type at 12 hours, when *IME1* was overexpressed (Fig 3C). This suggests that the second block in meiosis in *kar4*Δ/Δ is at least partially upstream of *NDT80* expression.

**Fig 3.**
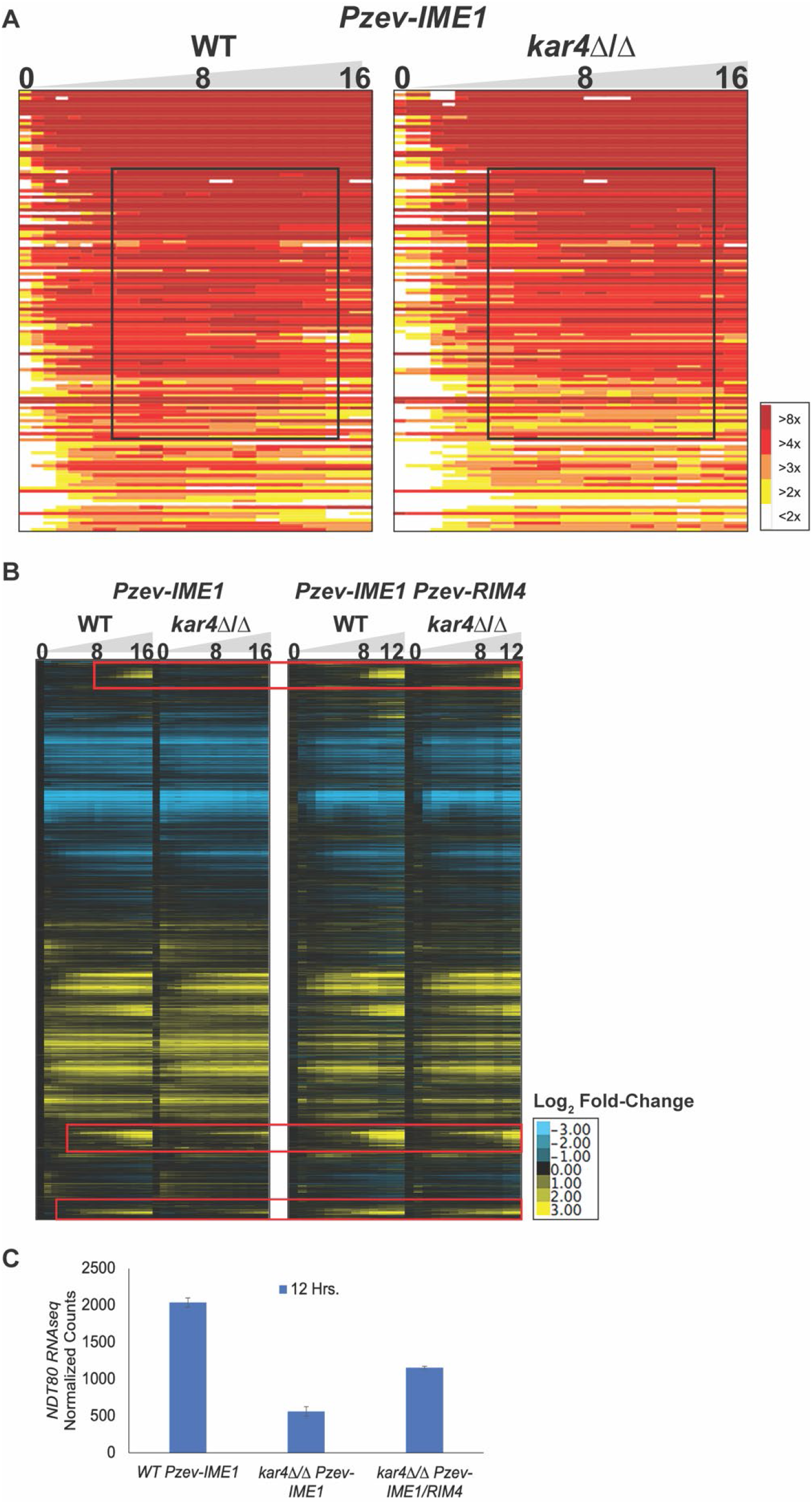
*IME1* and *RIM4* overexpression rescue the majority of the *kar4*Δ/Δ transcript level defect. (A) Heatmap of microarray data of Ime1p dependent genes after *IME1* overexpression in wild type and *kar4*Δ/Δ. Black box highlights the ability of *IME1* overexpression to rescue the defect in expression of these genes seen in *kar4*Δ/Δ without *IME1* overexpression. (B) Heatmap of microarray data in wild type and *kar4*Δ/Δ with *IME1* overexpressed (Left) and *IME1* and *RIM4* overexpressed (Right). Red boxes highlight the three gene clusters that show a defect in expression in *kar4*Δ/Δ after *IME1* overexpression but are rescued to some extent by additionally overexpressing *RIM4*. (C) *NDT80* RNA-seq normalized counts from wild type and *kar4*Δ/Δ with *IME1* overexpressed and from *kar4*Δ/Δ with *IME1* and *RIM4* overexpressed. Counts were normalized using the standard normalization method in DESeq2. Error bars represent standard deviation between two biological replicates.

### *RIM4* and *IME1* Co-Overexpression Suppresses the Late *kar4*Δ/Δ Transcript Abundance Defects

Co-overexpression of *RIM4* and *IME1* is necessary to fully suppress the *kar4*Δ/Δ meiotic defects. To understand the basis for suppression, we performed gene expression profiling on wild type and *kar4*Δ/Δ strains with both Pzev-*IME1* and Pzev-*RIM4*. First, the expression data showed that the dual induction system does overexpress both *IME1* and *RIM4*. Second, with both suppressor genes over-expressed, late gene expression was largely restored in the *kar4*Δ/Δ strain. Third, in wild type when Ime1p and Rim4p are co-overexpressed, the gene expression profile showed faster progression through the early meiotic transcriptional regime, relative to Ime1p expression alone (S Fig 1). However, in *kar4*Δ/Δ, an increase in the speed of progression through the early meiotic transcriptional regime is not as pronounced (S Fig 1) and there remained a moderate delay or reduction in the expression of *NDT80* and the *NDT80*-regulon in *kar4*Δ/Δ (Fig 3B, Fig 3C), relative to wild-type in which *IME1* and *RIM4* are overexpressed.

This delay most likely explains why *kar4*Δ/Δ strains do not sporulate as well as wild type when both *IME1* and *RIM4* are co-overexpressed. Nevertheless, the co-overexpression of both suppressor genes does restore sporulation to *kar4*Δ/Δ to a level comparable to what is observed in wild type S288c cells (see companion manuscript).

### Decreased Ime2p Expression is not Responsible for the Defects in Meiotic Progression

Rim4p was first identified as a positive regulator of early meiotic gene expression including the expression of the meiotic kinase, *IME2* (Soushko and Mitchell 2000, Deng and Saunders 2001). Previous work showed that *IME2* transcripts are methylated during meiosis (Bodi, Button et al. 2010, Schwartz, Agarwala et al. 2013). Accordingly, we asked if Kar4p also plays a role in regulating *IME2* expression. Ime2p levels were measured during meiosis using an epitope tagged Ime2p. In both wild type and *kar4*Δ/Δ cells, the level of Ime2p rose throughout the time course. Interestingly, we found no difference in the level of *IME2* transcript or Ime2p between wild type and *kar4*Δ/Δ across a time course of meiosis (S Fig 2). This suggests that Ime2p is not limiting the progression of *kar4*Δ/Δ cells through pre-meiotic DNA synthesis and recombination, consistent with the prior defect in Ime1p expression. However, it is possible that differences in Ime2p levels might arise later in meiosis as wild type continues to progress through the meiotic program and *kar4*Δ/Δ does not.

Because *IME1* overexpression allows progression past the initial meiotic block, we checked for defects in Ime2p expression that appear later in meiosis. Accordingly, we examined Ime2p levels in *kar4*Δ/Δ and wild type after overexpression of *IME1*, in the absence of *RIM4* overexpression. *IME2* transcript and Ime2p levels were reduced only 2-fold at 12 hours post *IME1* induction in *kar4*Δ/Δ compared to wild type (Fig 4A, Fig 4B). However, expression of *IME2* transcript and protein was similar to wild type at earlier time points and it was only the late burst of expression that was absent in *kar4*Δ/Δ (Fig 4A, Fig 4B). The deficit in *IME2* expression at 12 hours between wild type and *kar4*Δ/Δ could be due to the delay in *NDT80* expression in *kar4*Δ/Δ (Fig 3C) since mutants defective for *NDT80* expression also lose this late burst of Ime2p expression (Shin, Skokotas et al. 2010). Taken together, this suggests that defects in Ime2p expression are not solely responsible for the continued loss of sporulation in *kar4*Δ/Δ after *IME1* overexpression.

**Fig 4.**
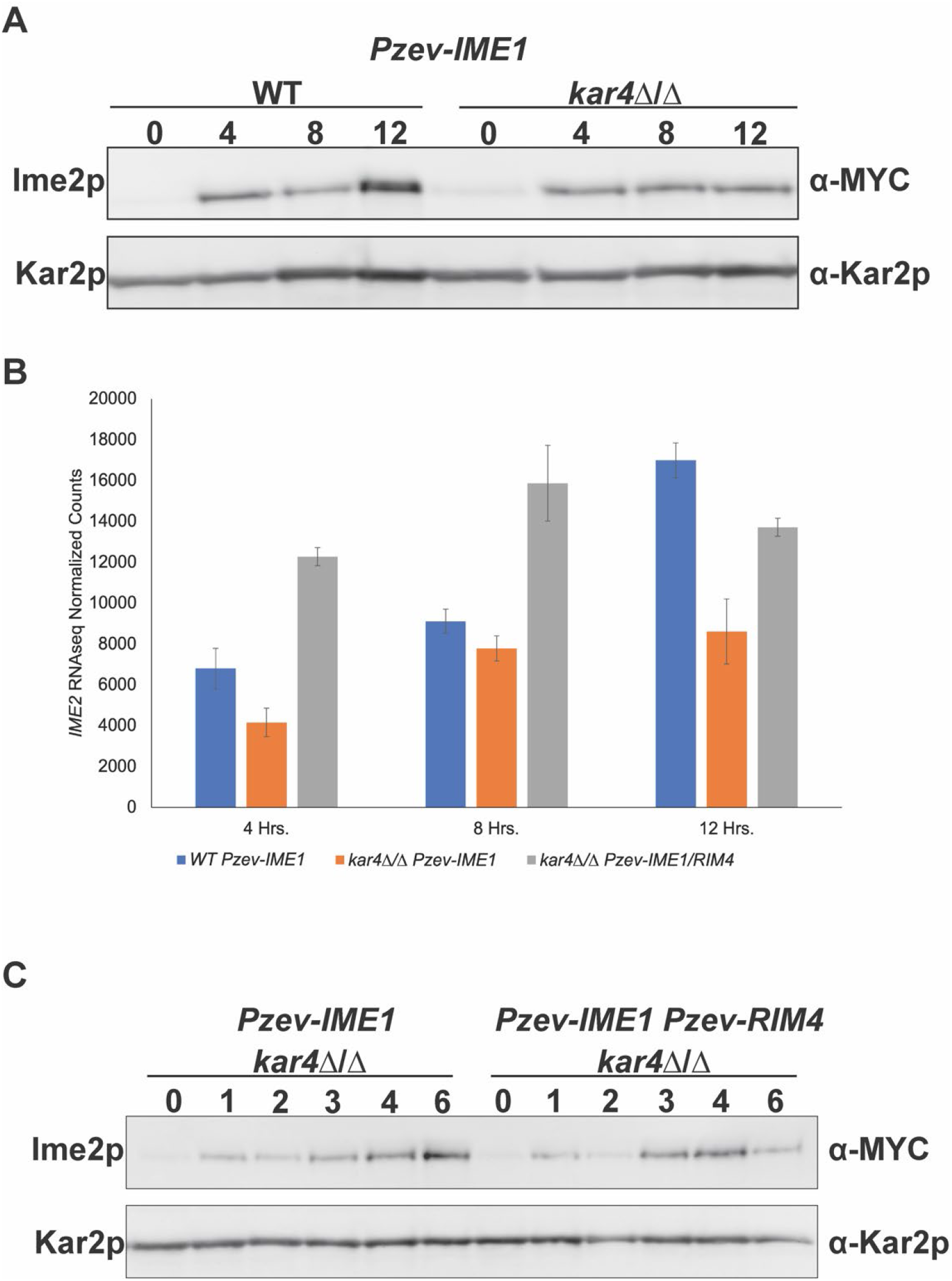
Defects in Ime2p expression are not responsible for block in meiotic progression. (A) Western blots of Ime2p-13MYC across a time course of meiosis in wild type and *kar4*Δ/Δ with *IME1* overexpressed. Kar2p is used as a loading control. (B) *IME2* RNA-seq normalized counts from wild type and *kar4*Δ/Δ with *IME1* overexpressed as well as *kar4*Δ/Δ with *IME1* and *RIM4* overexpressed. Counts were normalized using the standard normalization method in DESeq2. Error bars represent standard deviation between two biological replicates. (C) Western blots of Ime2p-13MYC across a time course of meiosis in *kar4*Δ/Δ with either *IME1* overexpressed or *IME1* and *RIM4* overexpressed. Kar2p is used as a loading control.

To determine the impact of *RIM4* overexpression on Ime2p levels, we assayed Ime2p in *kar4*Δ/Δ with both *IME1* and *RIM4* overexpressed. In this strain, *IME2* transcript levels peaked earlier than when only *IME1* is overexpressed (Fig 4B). Therefore, we looked at Ime2p at earlier time points in *kar4*Δ/Δ with either *IME1* overexpressed or *IME1* and *RIM4* co-overexpressed.

Ime2p levels peaked at 4 hours in the double overexpression strain and then begin to go down by 6 hours whereas Ime2p levels continue to increase in *kar4*Δ/Δ at these times with just *IME1* overexpressed (Fig 4C). Thus, the additional overexpression of *RIM4* sped up, but did not cause higher levels, of the expression of Ime2p in *kar4*Δ/Δ.

### Kar4p is Required for the Level of Multiple Meiotic Proteins

Given that Ime2p levels did not appear to be limiting the progression of *kar4*Δ/Δ after *IME1* overexpression, but that the second block is suppressed by a known translational regulator, we hypothesized that there may be critical regulatory proteins whose expression is impacted in *kar4*Δ/Δ. To identify candidate proteins, mass spectrometry was used to identify proteins whose levels are strongly dependent on Kar4p, but whose transcript levels are not. The P_Z3EV_-*IME1* strains were used for three reasons: first, they show greater meiotic synchrony; second, *IME1* overexpression suppresses the early transcript abundance defects in *kar4*Δ/Δ; and third, *IME1* overexpression alone can suppress the defect of a catalytic mutant of Ime4p, suggesting that defects that persist after *IME1* overexpression do not involve mRNA methylation. Because meiotic defects appear at a later stage in *IME1*-overexpressed cells, we examined proteins 8- and 12-hours post-*IME1* induction. Total protein samples were digested with trypsin, fractionated using ion exchange, and analyzed by liquid chromatography - mass spectrometry/mass spectrometry (LC-MS/MS).

Using LC-MS/MS, we were able to identify 4068 proteins expressed during meiosis. Spectral count data from the mass spectrometry was used to approximate relative protein levels. We used a cutoff of proteins reduced more than 2-fold in *kar4∆/∆* cells relative to wild type. At 8 hours, 432 proteins were present at less than 50% the levels in *kar4*Δ/Δ compared to wild type.

GO term analysis of these low proteins at 8 hours returned “meiotic cell cycle” as the sixth term with a P value of 7.74×10^−5^. At 12 hours, 318 proteins were present at less than 50% the levels in *kar4*Δ/Δ relative to wild type. GO term analysis of these low proteins at 12 hours returned “meiotic cell cycle” as the second term with a P value less than 0.001. Many proteins were low at both 8 and 12 hours in *kar4∆/∆* including Sps2p, Gas4p, Gmc2p, Mei5p, and Sae3p but there were also proteins including Ecm11p, Hed1p, Spo11p, and Rec8p that were expressed at or above wild type levels at 8 hours and then went down to lower than wild type levels at 12 hours.

To validate some of the candidate proteins identified by mass-spectrometry, we epitope-tagged two proteins of interest (Gas4p and Sps2p) and assayed their protein levels using western blotting and transcript levels with qPCR. We were not able to detect either Gas4p or Sps2p at 8 hours post *IME1* induction in either wild type or *kar4∆/∆*, which supports the mass spec data (Fig 5A). At 12 hours post *IME1* induction in wild type, both Gas4p and Sps2p were detectable, and this was accompanied by increased transcript levels (Fig 5A, Fig 5B). However, in *kar4∆/∆* at 12 hours there was still no detectable Gas4p or Sps2p. We determined that we could have detected 10-fold and 14-fold lower levels of Gas4p and Sps2p, respectively, relative to wild-type at 12 hours post *IME1* induction. Thus, the reductions in protein abundance were much larger than the 3-fold reduction observed at the transcript level (Fig 5A, Fig 5B). It is important to note that both *GAS4* and *SPS2* are under the control of Ndt80p, which is reduced 3-fold in *kar4∆/∆* with *IME1* overexpression (Fig 3C).

**Fig 5.**
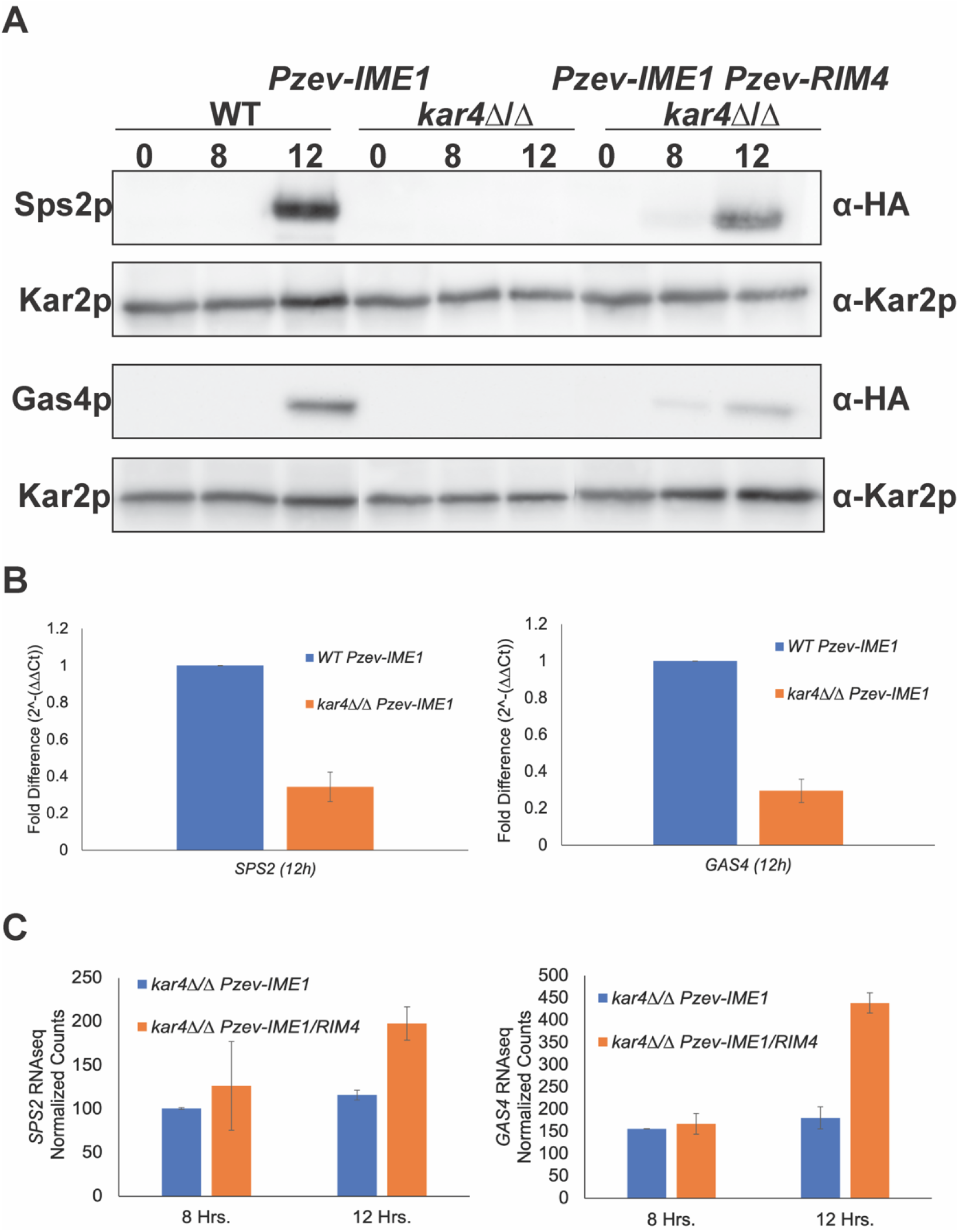
Kar4p is required for wild type levels of several meiotic proteins. (A) Western blots of Sps2p-3HA and Gas4p-3HA in wild type and *kar4*Δ/Δ with *IME1* overexpressed and in *kar4*Δ/Δ with *IME1* and *RIM4* overexpressed. Blots in the same row were exposed for the same length of time. Kar2p serves as a loading control. (B) qPCR measurements of the change in expression for SPS2 (Left) and GAS4 (Right) between wild type and *kar4*Δ/Δ with *IME1* overexpressed. Fold changes were calculated using the ΔΔCt method and *PGK1* was used as the normalizing gene. (C) SPS2 (Left) and GAS4 (Right) RNA-seq normalized counts from *kar4*Δ/Δ with either *IME1* overexpressed or *IME1* and *RIM4* overexpressed. Counts were normalized using the standard normalization method in DESeq2. Error bars represent standard deviation between two biological replicates.

We next wanted to determine how the additional overexpression of *RIM4* can suppress the later *kar4*Δ/Δ defect. Remarkably, upon *IME1* and *RIM4* overexpression, Gas4p and Sps2p become detectable at 8 hours in *kar4*Δ/Δ, but there was no change in transcript abundance relative to *kar4*Δ/Δ with only *IME1* overexpressed (Fig 5A, Fig 5C). At 12 hours, both proteins were strongly expressed, although not at quite as high levels as in wild type with only *IME1* overexpressed. The increase was accompanied by only a 2-fold increase in transcript abundance. Thus, the relatively small changes in transcript levels were accompanied by at least 10- to 20- fold increases in the levels of Gas4p and Sps2p, respectively, suggesting that the overexpression of *RIM4* is functioning to enhance translation of these two proteins. This finding point to a potential mechanism of the early positive acting meiotic function of Rim4p.

### Defects in the Expression of Recombination Proteins Persist After *IME1* Overexpression in *kar4*Δ/Δ

Although the experiments described above validate the mass spectrometry and microarray/RNA-seq data, it is likely that the second block in meiosis is caused by reduced levels of proteins that act earlier than Sps2p and Gas4p. From the mass spectrometry data, key recombination proteins were found to be mis-regulated in *kar4∆/∆* including Mei5p, Sae3p, Gmc2p, and Ecm11p. All four proteins are highly reduced at 12 hours in *kar4∆/∆*, but we see little to no impact on their transcript levels (Fig 6A-D) suggesting that Kar4p could be positively regulating the translation of these proteins as opposed to their transcript abundance.

**Fig 6.**
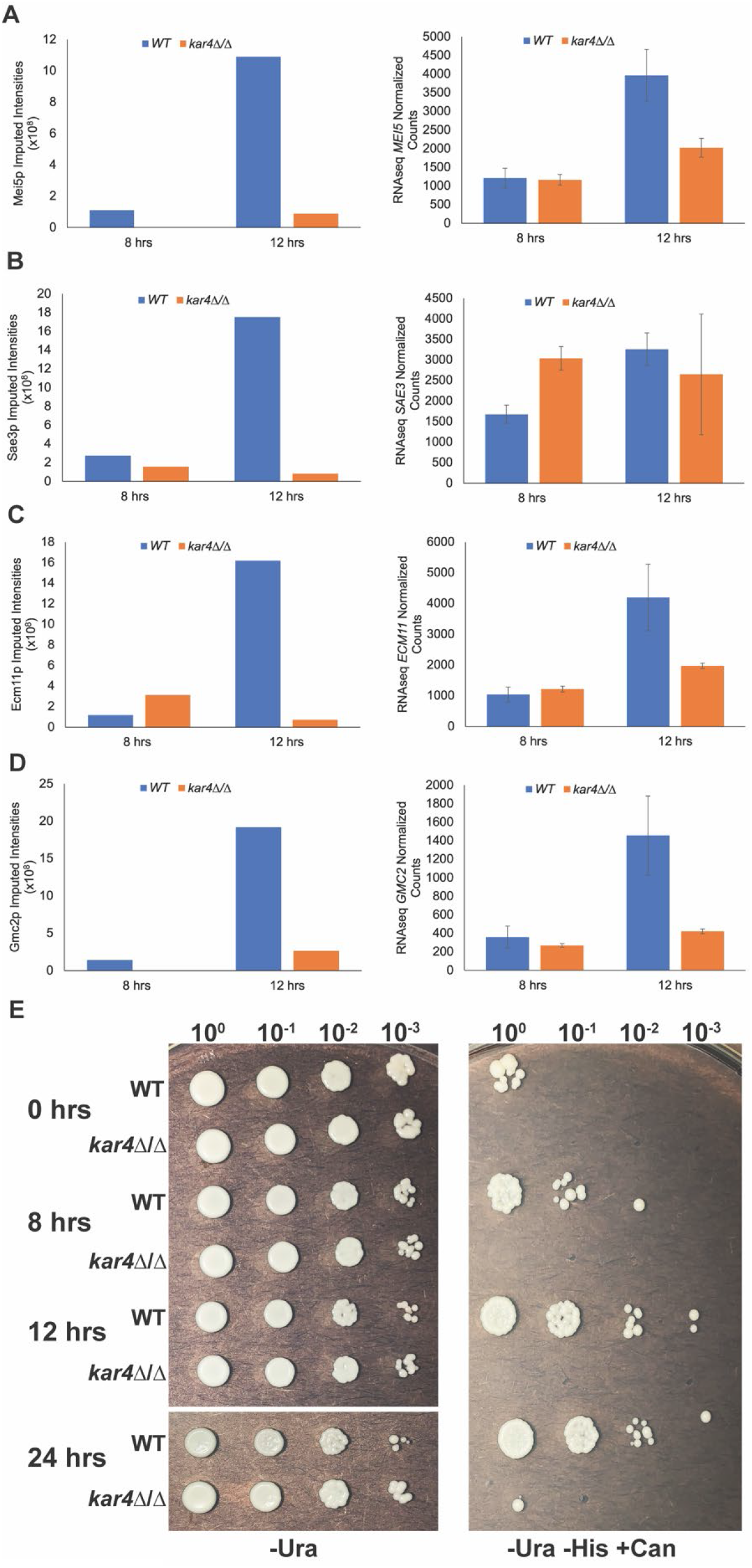
Kar4p is required for the expression of key recombination proteins. Protein levels measured by mass spectrometry (left) and RNA-seq normalized counts (right) of (A) *MEI5*, (B) *SAE3*, (C) *ECM11*, and (D) *GMC2*. Error bars represent standard deviation of two biological replicates. Normalized counts were calculated using DESeq2. (E) Screen for recombination described in a companion manuscript. Wild type and *kar4*Δ/Δ both carry *IME1* on a high-copy number plasmid also carrying the *URA3* gene. Growth on SC-Ura (left) was used to assay growth and maintenance of the plasmid. Growth on SC-Ura -His +can (right) was used to select for recombination events. Spots are 10-fold serial dilutions of a starting concentration of 1 OD unit of cells for each strain at each time point (0, 8, 12, and 24 hours post movement into sporulation media).

Interestingly, we also saw lower levels of some of these proteins (Mei5p, Gmc2p, and Sae3p) at the 8-hour time point (Fig 6A, Fig 6B, Fig 6D), which would suggest that defects in recombination should be present in *kar4∆/∆* even at this relatively early time point. To address this, we examined the timing of meiotic recombination in wild type and *kar4*∆/∆ cells carrying a high-copy number plasmid containing *IME1* (see companion manuscript). In wild type cells, cells that have undergone recombination begin to appear at 8 hours after transfer to sporulation conditions, increasing greater than 10-fold over the next 4 hours (Fig 6E). As predicted, in cells lacking Kar4p, cells that had undergone recombination were not observed even after 24 hours (Fig 6E). The initial screen for high-copy suppressors that identified *IME1* allowed *kar4∆/∆* cells to sporulate for up to 4 days, indicating that these cells do eventually undergo recombination, but the timing is delayed (see companion manuscript). Together, these data suggest that Kar4p positively regulates the translation of proteins important for meiotic recombination including Mei5p, Sae3p, Gmc2p, and Ecm11p and loss of this regulation impairs the efficiency of meiotic recombination. The persistence of the expression defects, despite the ability of *IME1* overexpression to suppress a catalytically dead mutant of Ime4p (see companion manuscript), implies that they reflect a loss of a function of Kar4p separate from mRNA methylation.

## Discussion

Through analyzing both the transcriptome and proteome of *kar4*Δ/Δ mutants during meiosis we now have a better understanding of the molecular underpinnings underlying the *kar4*Δ/Δ meiotic defects, and how overexpression of *IME1* and *RIM4* suppress the defects. Establishing a more defined set of Ime1p-dependent genes demonstrated the impact of loss of Kar4p on the *IME1* regulon. Overexpression of *IME1* revealed that a later block in meiosis is at least partially upstream of *NDT80* expression. *RIM4* overexpression appears to rescue this later defect by impacting the translation of transcripts as opposed to facilitating increased gene expression.

Microarray and RNA-seq data showed that *kar4*Δ/Δ mutants have a wild type transcriptional profile with two exceptions: first, they have an early defect in the transcript level of *IME1*, as well as a subset of Ime1p-dependent genes that are not immediately activated by the early burst of *IME1* expression that is independent of mRNA methylation. The early defect is suppressed by the overexpression of *IME1*. Second, a late defect is not suppressed by *IME1* overexpression alone, but is suppressed by the additional overexpression of *RIM4*. However, the late transcriptional defect is most likely an indirect effect of a prior arrest point of *kar4*Δ/Δ mutants suppressed by *IME1* overexpression. The late genes that are impacted are downstream of Ndt80p, but the block in *kar4*Δ/Δ after *IME1* overexpression is upstream of Ndt80p expression. Thus, the low levels of the late genes are most likely due to the loss of expression of Ndt80p.

The early defect in *IME1* expression and the expression of Ime1p dependent genes support findings in a companion manuscript that show lower levels of Ime1p and *IME1* transcript in *kar4*Δ/Δ during meiosis. Therefore, the overexpression of *IME1* bypasses the requirement of Kar4p in regulating the expression of *IME1* and its targets. The later block in gene expression after *IME1* overexpression suggests that Kar4p is required at another step that impacts the induction of Ndt80p.

Given that the overexpression of the translational regulator *RIM4* is required to suppress the later *kar4*Δ/Δ meiotic defect, we hypothesize that Kar4p may also have a role in translational regulation during meiosis. Rim4p’s role in promoting the expression of Ime2p made it a potential candidate for Kar4p translational regulation. With *IME1* under its own promoter, we saw no difference in the expression of *IME2* transcript or Ime2p between wild type and *kar4*Δ/Δ. Thus, it is unlikely that Ime2p is limiting the progression of *kar4*Δ/Δ cells early in meiosis. However, there is a defect in both *IME2* transcript and Ime2p levels that is revealed when *IME1* is overexpressed. Ime2p is required for the activation of the mid-meiotic transcription factor Ndt80p, which initiates the expression of genes required for the completion of meiotic recombination and prophase I exit as well as the meiotic divisions and spore maturation. Low Ime2p levels were rescued in *kar4*Δ/Δ when both *IME1* and *RIM4* were overexpressed. In addition, the overexpression of both *IME1* and *RIM4* together significantly sped up the time to peak expression of Ime2p in *kar4*Δ/Δ.

We initially screened Ime2p levels because Rim4p is required for Ime2p expression, but an unbiased mass spectrometry approach identified many proteins that are reduced in *kar4*Δ/Δ at both 8 and 12 hours post induction of *IME1* expression. Many of these proteins were important for meiotic recombination including Mei5p, Gmc2p, Sae3p, and Ecm11p. These proteins showed little or no difference in transcript levels between wild type and *kar4*Δ/Δ at 8 and 12 hours, implying that the reduced protein levels at 8 and 12 hours in *kar4*Δ/Δ is likely due to defects in translation as opposed to transcript abundance. Consistent with the fact that many of the impacted proteins are involved in recombination, we found that recombination was delayed in *kar4*Δ/Δ, even with *IME1* overexpressed. Defects in recombination would activate the meiotic recombination checkpoint mediated by Mek1p (Chen, Gaglione et al. 2018), which acts antagonistically to Ime2p to block the activation of Ndt80p. Activation of the *MEK1* checkpoint could explain the defects in *NDT80* expression in *kar4*Δ/Δ, as well as in Ime2p expression; Ndt80p also induces *IME2* expression, resulting in Ime2p and Ndt80p levels peaking at similar times later in meiosis (Shin, Skokotas et al. 2010). Because overexpression of *IME1* suppresses the meiotic defect of a catalytic mutant of Ime4p, we hypothesize that the defect in protein expression involves the proposed non-catalytic function of the methyl-transferase complex.

Consistent with this, recent work has shown that the ortholog of Ime4p in mammals, METTL3, positively regulates the translation of transcripts in an m^6^A-independent manner by interacting with PABP and cap-binding factors (Lin, Choe et al. 2016, Wei, Huo et al. 2022). Given that METTL3 and METTL14 work together to bind mRNA, it is likely that METTL14, Kar4p’s ortholog, is also involved in this function. However, due to the nature of the arrest after *IME1* overexpression, we cannot determine if mRNA methylation is also important for events downstream of Ndt80p. Future work will determine if Kar4p and other members of the complex are regulating the translation of transcripts in yeast in a similar manner and if mRNA methylation plays a role in regulating later steps of meiosis.

The co-overexpression of *IME1* and *RIM4* partially rescued the protein level defects. Rim4p is best known as a repressor of translation that functions as an aggregate to sequester mRNAs from the translational machinery (Berchowitz, Kabachinski et al. 2015). However, Rim4p was first identified as a positive regulator of *IME2* (Deng and Saunders 2001). No further work was done to explore the positive regulatory role of Rim4p. The fact that overexpression of *RIM4* rescues the *kar4*Δ/Δ translational defect suggests that Rim4p acts directly as an enhancer of translation or that the overexpression facilitates Rim4p aggregates to expand their regulon and sequester a negative regulator that is not normally bound by the aggregates. Work on an analogous protein to Rim4p in mammals, “Deleted in Azoospermia Like” (DAZL), has shown that DAZL exists as both a monomer and an aggregate. The aggregate acts to block translation and the monomeric form promotes translation through interactions with PABP (Collier, Gorgoni et al. 2005, Berchowitz, Kabachinski et al. 2015, Laureau, Dyatel et al. 2021). Rim4p monomers are seen early in meiosis suggesting that it is the monomeric form of Rim4p that is important for its role in meiotic entry and these monomers may act to positively regulate translation in a similar manner to DAZL monomers. An interaction between Kar4p and Rim4p has not been detected, which suggests that the suppression bypasses the requirement for Kar4p.

Taken together, these data point to Kar4p being a key player in the post-transcriptional/translational regulation of meiosis. In support of Kar4p’s role in mRNA methylation, we see that expression of both *IME1*, and Ime1p-dependent genes are lower in *kar4*Δ/Δ during meiosis. These results indicate that Kar4p is required upstream of *IME1* expression. That *IME1* overexpression bypasses the requirement for Kar4p in meiotic entry, but cells remain blocked before the induction of *NDT80* expression, suggests that Kar4p is required at another step during meiosis. That the additional overexpression of *RIM4* allows *kar4*Δ/Δ cells to complete sporulation points to Kar4p playing a role in translational regulation that is important upstream (and possibly downstream) of *NDT80* expression. Interestingly, deletions of other methylation complex members (Ime4p and Mum2p) can also be made to sporulate after overexpression of both *IME1* and *RIM4* (see companion manuscript). Future work will determine whether the regulation of these protein levels is similar to the non-catalytic functions described in other eukaryotes. Understanding the non-catalytic function of Kar4p may also provide insights into how Rim4p functions positively in meiosis. These findings position Kar4p as a key regulator of meiosis at multiple levels and further experimentation will seek to determine how exactly this intrepid protein is carrying out that regulation.

## Materials and Methods

### Sporulation

Cultures were grown overnight at 30 °C in YPD (yeast nitrogen base (1% w/v), peptone (2% w/v), and 2% glucose), back diluted into YPA (yeast nitrogen base (1% w/v), peptone (2% w/v), and potassium acetate (1% w/v), and allowed to grow for 16-18 hours before being transferred into 1% (w/v) potassium acetate sporulation media supplemented with histidine, uracil, and leucine at a concentration of 0.5 OD_600_ unit/ml. Sporulating cells were cultured at 26°C for various amounts of time depending on the nature of the experiment. For experiments involving overexpression, 1 µM of β-estradiol was added to cultures once they were moved into the 1% potassium acetate media.

### RNA Preparation

Cells from sporulation cultures were collected by vacuum filtration on nitrocellulose filters and flash frozen with liquid nitrogen. Samples were stored at -80°C until extracted for RNA. RNA was extracted by acid-phenol method. Lysis buffer was added and then vigorously vortexed. Following lysis, phenol saturated with 0.1 M citrate (Sigma-Aldrich, P4682) was added. Lysates were incubated for 30 minutes at 65°C, vortexing every 5 minutes. Lysates were chilled on ice and then spun. Supernatant was added to a heavy phase lock tube along with chloroform. After light mixing, the samples were centrifuged. The aqueous layer was moved to a new tube containing sodium acetate. The samples were washed with ice-cold 100% ethanol and left to incubate at -20°C for 30 minutes to 16 hours. Following the incubation, ethanol-precipitated samples were spun at full speed for 5 minutes to pellet the RNA. The pellets were washed with ice cold 70% ethanol and pulse spun. Remaining alcohol was aspirated, and the RNA samples were resuspended in 100 μl of water. RNA samples were cleaned up using the Qiagen RNeasey kit (Qiagen 74106) and then quantified using a Nanodrop.

### Gene Expression Microarrays

Microarray analysis was performed as described by Brauer et al. 2008 with slight modifications: RNA samples were handled in an ozone-free environment during the labeling process and labeling was performed using the Quick Amp labeling kit (5190-0447)) according to a modified labeling protocol. Reference RNA was labeled with Cy3-CTP (NEL580) and experimental samples were labeled with Cy5-CTP (NEL581). Labeled cRNA samples were cleaned up using the Qiagen RNeasy Cleanup kit protocol with an additional wash step, then quantified using a Nanodrop. Labeled cRNA was fragmented and allowed to hybridize to agilent microarray slides (8×15k, AMADID:017566) for 17 hours at 65°C and 20 RPM. After hybridization, slides were washed successively with wash buffer 1 for 1 minute, washer buffer 2 for 1 minute, and acetonitrile for 30 seconds. Slides were scanned using the Agilent High-Resolution Microarray Scanner. After scanning, Feature Extraction software was used to map spots to the specific genes. Resulting microarray intensity data were submitted to the PUMA Database (http://puma.princeton.edu) for archiving and analysis.

### Microarray Analysis

Sample and reference channel intensities were first floored to a value of 350. Once log2 ratios were computed between samples and reference, the data were time-zero transformed. Data were hierarchically clustered in the Cluster 3.0 software package with average linkage using the Pearson correlation distance as the metric of similarity between genes. Gene Ontology terms were determined using YEASTTRACT.

### RNA-seq

Cells were induced to sporulate as described above and samples were taken at the indicated time points. Cells were lysed using bead beating and the lysis buffer included in the Qiagen RNeasy Kit. After lysis, samples were cleared by centrifugation and RNA was purified using the Qiagen RNeasy kit with on-column DNase treatment. RNA samples were then sent to Novogene Corporation for library prep and mRNA sequencing using an Illumina based platform (PE150). Resulting data was analyzed using the open access Galaxy platform (Afgan, Baker et al. 2018). Reads were first mapped to the yeast genome using BWA-MEM and then counted using htseq-count. Differential expression analysis was conducted using DESeq2 and heat maps were made using Cluster 3.0 and Java Tree View.

### qPCR

Cells were induced to sporulate as described above and samples were taken at the indicated time points. RNA was harvested as described in the section on RNA-seq. cDNA libraries were constructed using the High-Capacity cDNA Reverse Transcription kit (Applied Biosystems) with 10 µl of the total RNA sample. The concentration of the resulting cDNA was measured using a nanodrop. qPCR reactions were set up using Power SYBR Green PCR Master Mix (Applied Biosystems) with 50 ng of total RNA. The reactions were run on a CFX96 Real-Time System (BioRad) with reaction settings exactly as described in the master mix instructions with the only change being the addition of a melt curve at the end of the program. Results were analyzed using CFX Maestro. Primer sequences were as follows: *PGK1* Forward 5’-CTCACTCTTCTATGGTCGCTTTC-3’, *PGK1* Reverse 5’-AATGGTCTGGTTGGGTTCTC-3’, *GAS4* Forward 5’-GACCTGGAAGGAGAAGAAGAACAAG-3’, *GAS4* Reverse 5’-ACAATGGGCCGGAAATAGAG-3’, *SPS2* Forward 5’-GCCGGTCGTTCGATCATAA-3’, *SPS2* Reverse 5’-CATTGTCAGTTTCCTGCTTTCC-3’.

### Protein Extraction by Alkaline Lysis

Optical density was measured and a total of 6 OD_600_ units was collected for each time point. Cells were pelleted and stored at -80°C. Cell lysates were prepared by adding 150 μl lysis buffer (1.85 M NaOH, 1/100 B-ME, 1/50 protease inhibitors) followed by a 10-minute incubation on ice. After incubation, 150 μl of 50% TCA (Sigma-Aldrich T9159) was added.

After a 10-minute incubation at 4°C, samples were spun for 15 seconds at 15000 RPM and supernatant was aspirated. The remaining pellet was washed with one ml acetone, briefly spun, and acetone aspirated. 100 μl of 2x sample buffer (ThermoFisher NP0007) with 10% B-ME were added to the protein pellets, mixed well, and boiled for 5 minutes.

### Immunoblotting

For each lane, 10 μl of protein extracts were added to 8% Bis-Tris acrylamide gels. Protein ladder (Precision Plus Protein Standard from Bio Rad, 1610374) was used for determining band sizes. Electrophoresis was run at 60 volts for 30 minutes through the stacking gel and at 150 volts until the samples moved through the resolving gel. Gels were transferred to PVDF membranes using a semi-dry transfer apparatus (TransBlot SD BioRad) and a standard Tris-Glycine transfer buffer without methanol at 16 volts for 36 minutes. Membranes were blocked with 10% milk for 30 minutes, followed by primary antibody (anti-MYC 1:1000 (9E10) and anti-HA 1:1000 (12CA5)) in 0.1% TBST for 1 hour with rocking at room temperature. Membranes were washed three times for 10 minutes with TBST. Membranes were incubated with secondary antibody (Donkey anti-mouse (Jackson ImmunoResearch) IgG 1:10,000) in 1% milk with rocking for 30 minutes. Membranes were then washed three times for 10 minutes with TBST. Immobilon Western HRP substrate (Millipore) was added and incubated for 5 minutes before being imaged using the G-Box from SynGene. Densitometry was conducted using ImageJ. All westerns were run at least twice with each run being a unique biological replicate.

### Trypsin Digest and 8-Step Fractionation Mass Spectrometry

For each time point, the optical density was measured and a total of 30 OD_600_ was aliquoted. Samples were pelleted, washed with water, and flash frozen by liquid nitrogen. Frozen yeast pellets were resuspended in lysis buffer (6M guanidium hydrochloride, 10mM TCEP, 40mM CAA, 100mM Tris pH 8.5). Cells were lysed by sonication using 5x 30s pulses with 1min rest in ice between pulses. Samples were then heated to 95°C for 15 min, and allowed to cool in the dark for 30 min. Samples were then centrifuged, and lysate removed to a fresh tube. Lysate was then diluted 1:3 with digestion buffer (10% CAN, 25mM Tris pH 8.5) containing LysC (1:50) and incubated at 37°C for 3 hours. Samples were then further diluted to 1:10 with digestion buffer containing Trypsin (1:100) and incubated O/N at 37°C. TFA was added to 1% final. Samples were then centrifuged, and digested lysate removed to a new tube. Samples were desalted on C18 cartridges (Oasis, Waters) as per manufacturer protocol. Dried down peptide samples were then fractioned using High pH Reversed-Phase peptide fraction kit (Pierce) into 8 fractions using manufacturer’s instructions. Fractions were dried completely in a speedvac and resuspended with 20μl of 0.1% formic acid pH 3. 5ul was injected per run using an Easy-nLC 1000 UPLC system. Samples were loaded directly onto a 45cm long 75μm inner diameter nano capillary column packed with 1.9μm C18-AQ (Dr. Maisch, Germany) mated to metal emitter in-line with a Q-Exactive (Thermo Scientific, USA). The mass spectrometer was operated in data dependent mode with the 700,00 resolution MS1 scan (400-1800 m/z), AGC target of 1e6 and max fill time of 60ms. The top 15 most intense ions were selected (2.0 m/z isolation window) for fragmentation (28 NCE) with a resolution of 17,500, AGC target 2e4 and max fill time of 60ms. Dynamic exclusion list was evoked to exclude previously sequenced peptides for 120s if sequenced within the last 10s.

Raw files were searched with MaxQuant (ver 1.5.3.28) (Cox & Mann 2008), using default settings for LFQ data. Carbamidomethylation of cysteine was used as fixed modification, oxidation of methionine, and acetylation of protein N-termini were specified as dynamic modifications. Trypsin digestion with a maximum of 2 missed cleavages were allowed. Files were searched against the yeast SGD database download 13 Jan 2015 and supplemented with common contaminants. Results were imported into the Perseus (Tyanova et al. 2016) workflow for data trimming and imputation. Final data were exported as a table.

### Recombination Assay

Diploid wild type and *kar4*Δ/Δ strains carrying a P_*MFA1*_*-HIS3* cassette integrated in place of the *CAN1* gene in the *MAT*α parent were transformed with a high-copy number plasmid containing *IME1* and the *URA3* gene as a selectable marker. Strains were induced to sporulate as described above and samples were taken across a time course of meiosis (0, 8, 12, and 24 hours post movement into sporulation media). At each time point, 1 OD unit of cells was removed and serially diluted 10-fold three times. Four microliters of each serial dilution (10^0^, 10^−1^, 10^−2^, and 10^−3^) were plated on either SC-Ura (growth plate) or SC-Ura -His +can (recombination selection plate). Plates were allowed to grow for 2-3 days at 30°C before pictures were taken.

### Data Availability

Source data for the microarray and RNA-seq experiments can be found using GEO ascension numbers GSE220125 and GSE221451, respectively.

## Acknowledgements

We Anne Rosenwald for helpful feedback on this project and manuscript, and members of the Rose lab, especially Abigail Sporer who initiated this project and May Husseini for making media and reagents. We also thank Patrick Gibney and Scott McIsaac for feedback on this project and the construction of the estradiol inducible promoters. We especially thank David Botstein and Kara Dolinski for financial support of the microarray experiments. Additionally, we would like to thank the staff at the Princeton core facilities, especially Tharan Srikumar for his expert Mass Spectrometry, and John Matese, for help with Genomic databases. This work was supported by NIH grants GM037739 and GM126998 to MDR.

---

**S Fig 1.**
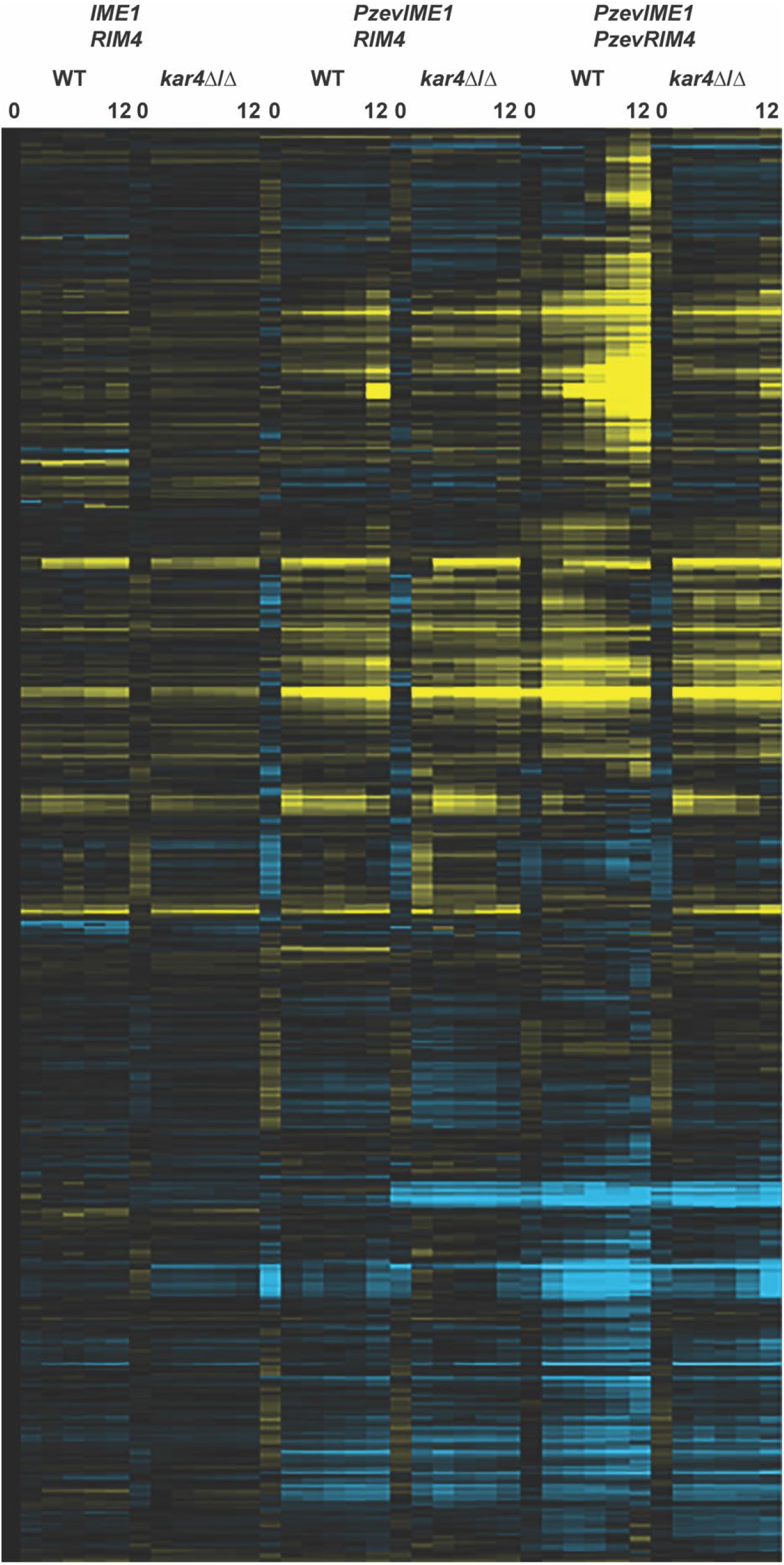
RNA-seq derived meiotic transcriptome of *kar4*Δ/Δ across overexpression conditions. Heatmap of RNA-seq data across a time course of meiosis (0, 2, 4, 6, 8, and 12 hours) in wild type and *kar4*Δ/Δ with either *pIME1/pRIM4, Pzev-IME1/pRIM4*, or *Pzev-IME1/Pzev-RIM4*. Expression was normalized to wild type pre-induction of sporulation (t=0). Genes were clustered in Cluster3.0 and the heatmaps were constructed with Java TreeView. Note that genes are clustered differently from Figure 1.

**S Fig 2.**
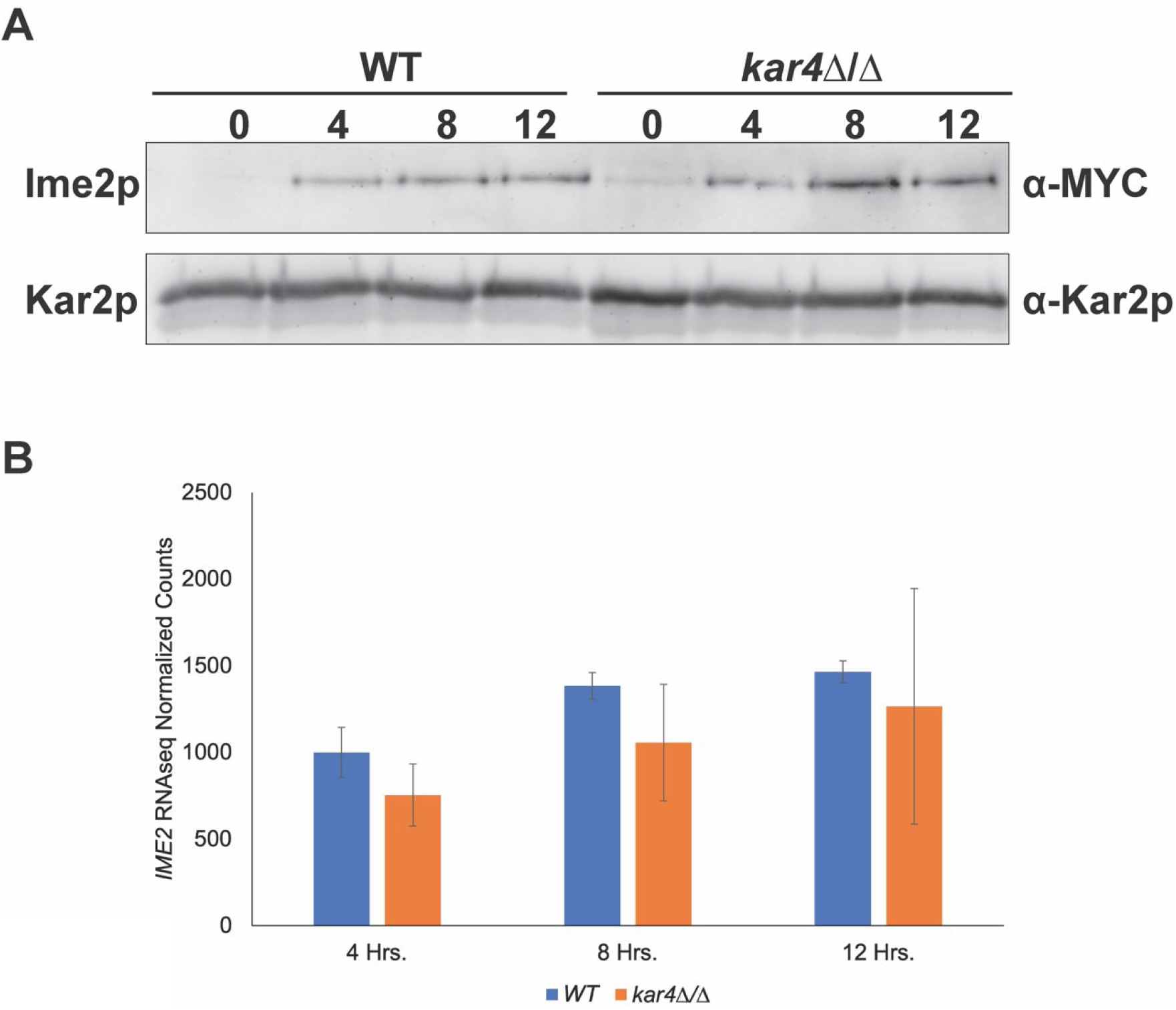
Ime2p levels are not impacted in *kar4*Δ/Δ under endogenous expression conditions. (A) Western blots of Ime2p-13MYC across a meiotic time course in wild type, *kar4*Δ/Δ, and *kar4*Δ/Δ with *IME1* overexpressed. Kar2p is used as a loading control. (B) *IME2* RNA-seq normalized counts from wild type, *kar4*Δ/Δ, and *kar4*Δ/Δ with *IME1* overexpressed. Counts were normalized using the standard normalization method in DESeq2. Error bars represent standard deviation between two biological replicates.

**Table S1.** Strains used for this study. All auxotrophic markers are standard BY alleles.

| Strain Name | Genotype | Strain |
| --- | --- | --- |
| MY 10128 | *leu2 his3 ura3 met15 kar4::KANMX* | S288c |
| MY 11297 | *ura3 leu2 his3 lys2 can1::LEU2+-MFA1pr-HIS3 kar4::KANMX* | S288c |
| MY 8092 | *leu2 his3 ura3 met15* | S288c |
| MY 16616 | *his3/" leu2/leu2::pACT1-Z3EV-NATMX +/lys2 met15/+ ura3/"* | S288c |
| MY 16533 | *HPHMX::PZ3EV-IME1/+ HPHMX::PZ3EV-RIM4/+ leu2::act1pr-Z3EV-NATMX/" ura3/"* | S288c |
| MY 16534 | *HPHMX::PZ3EV-IME1/+ leu2::act1pr-Z3EV-NATMX/leu2 ura3/" his3/+ lys2/+* | S288c |
| MY 16617 | *his3/" leu2/leu2::pACT1-Z3EV-NATMX +/lys2 met15/+ ura3/" kar4::KANMX/"* | S288c |
| MY 16531 | *HPHMX::PZ3EV-IME1/+ leu2::act1pr-Z3EV-NATMX/leu2 ura3/" his3/+ met15/+ kar4::KANMX/kar4::URA3* | S288c |
| MY 16536 | *HPHMX::PZ3EV-IME1/+ HPHMX::PZ3EV-RIM4/+ leu2::act1pr-Z3EV-NATMX/" ura3/" kar4::LEU2/kar4::URA3* | S288c |
| MY 16208 | *HPHMX::PZ3EV-IME1/+ leu2::act1pr-Z3EV-NATMX/leu2 ura3/" his3/+ met15/+ kar4::KANMX/+ GAS4-3HA::HIS3/+* | S288c |
| MY 16209 | *HPHMX::PZ3EV-IME1/+ leu2::act1pr-Z3EV-NATMX/leu2 ura3/" his3/+ met15/+ kar4::KANMX/+ SPS2-3HA::HIS3/+* | S288c |
| MY 16210 | *HPHMX::PZ3EV-IME1/+ leu2::act1pr-Z3EV-NATMX/leu2 ura3/" his3/+ met15/+ kar4::KANMX/kar4::URA3 GAS4-3HA::HIS3/+* | S288c |
| MY 16211 | *HPHMX::PZ3EV-IME1/+ leu2::act1pr-Z3EV-NATMX/leu2 ura3/" his3/+ met15/+ kar4::KANMX/kar4::URA3 SPS2-3HA::HIS3/+* | S288c |
| MY 16216 | *HPHMX::PZ3EV-IME1/+ HPHMX::PZ3EV-RIM4/+ leu2::act1pr-Z3EV-NATMX/" ura3/" kar4::KANMX/" GAS4-3HA::HIS3/+* | S288c |
| MY 16217 | *HPHMX::PZ3EV-IME1/+ HPHMX::PZ3EV-RIM4/+ leu2::act1pr-Z3EV-NATMX/" ura3/" kar4::KANMX/" SPS2-3HA::HIS3/+* | S288c |
| MY 16572 | *his3/" leu2/" ura3/" lys2/+ met15/+ IME2-13MYC::KANMX/+* | S288c |
| MY 16573 | *his3/" leu2/" ura3/" lys2/+ met15/+ IME2-13MYC::KANMX/+ kar4::HPHMX/"* | S288c |
| MY 16604 | *IME2-13MYC::KANMX/+ kar4::HPHMX/kar4::URA3 ura3/" his3/+ leu2/leu2::act1pr-Z3EV-NATMX HPHMX::PZ3EV-IME1/+ met15/+* | S288c |
| MY 16608 | *IME2-13MYC::KANMX/+ ura3/" his3/+ leu2/leu2::act1pr-Z3EV-NATMX HPHMX::PZ3EV-IME1/+ met15/+* | S288c |
| MY 16612 | *HPHMX::PZ3EV-IME1/+ HPHMX::PZ3EV-RIM4/+ kar4::URA3/kar4::LEU2 his3/+ leu2/leu2::act1pr-Z3EV-NATMX IME2-13MYC::KANMX/+ ura3/"* | S288c |

**Table S2.** Plasmids used in this paper.

| Strain | Markers | Source |
| --- | --- | --- |
| pMR<br>6895 | *2μ IME1 URA3 Amp* |  |
| pRB<br>3483 | *hphMx-Pz3ev CEN Amp* | McIsaac et al. 2014 |

## References

Afgan, E., D. Baker, B. Batut, M. van den Beek, D. Bouvier, M. Cech, J. Chilton, D. Clements, N. Coraor, B. A. Gruning, A. Guerler, J. Hillman-Jackson, S. Hiltemann, V. Jalili, H. Rasche, N. Soranzo, J. Goecks, J. Taylor, A. Nekrutenko and D. Blankenberg (2018). “The Galaxy platform for accessible, reproducible and collaborative biomedical analyses: 2018 update.” Nucleic Acids Res 46(W1): W537–W544.

Agarwala, S. D., H. G. Blitzblau, A. Hochwagen and G. R. Fink (2012). “RNA methylation by the MIS complex regulates a cell fate decision in yeast.” PLoS Genet 8(6): e1002732.

Ahmed, N. T., D. Bungard, M. E. Shin, M. Moore and E. Winter (2009). “The Ime2 protein kinase enhances the disassociation of the Sum1 repressor from middle meiotic promoters.” Mol Cell Biol 29(16): 4352–4362.

Berchowitz, L. E., A. S. Gajadhar, F. J. van Werven, A. A. De Rosa, M. L. Samoylova, G. A. Brar, Y. Xu, C. Xiao, B. Futcher, J. S. Weissman, F. M. White and A. Amon (2013). “A developmentally regulated translational control pathway establishes the meiotic chromosome segregation pattern.” Genes Dev 27(19): 2147–2163.

Berchowitz, L. E., G. Kabachinski, M. R. Walker, T. M. Carlile, W. V. Gilbert, T. U. Schwartz and A. Amon (2015). “Regulated Formation of an Amyloid-like Translational Repressor Governs Gametogenesis.” Cell 163(2): 406–418.

Bodi, Z., A. Bottley, N. Archer, S. T. May and R. G. Fray (2015). “Yeast m6A Methylated mRNAs Are Enriched on Translating Ribosomes during Meiosis, and under Rapamycin Treatment.” PLoS One 10(7): e0132090.

Bodi, Z., J. D. Button, D. Grierson and R. G. Fray (2010). “Yeast targets for mRNA methylation.” Nucleic acids research 38(16): 5327–5335.

Bowdish, K. S. and A. P. Mitchell (1993). “Bipartite structure of an early meiotic upstream activation sequence from Saccharomyces cerevisiae.” Mol Cell Biol 13(4): 2172–2181.

Bujnicki, J. M., M. Feder, M. Radlinska and R. M. Blumenthal (2002). “Structure prediction and phylogenetic analysis of a functionally diverse family of proteins homologous to the MT-A70 subunit of the human mRNA:m(6)A methyltransferase.” Journal of molecular evolution 55(4): 431–444.

Bushkin, G. G., D. Pincus, J. T. Morgan, K. Richardson, C. Lewis, S. H. Chan, D. P. Bartel and G. R. Fink (2019). “m(6)A modification of a 3’ UTR site reduces RME1 mRNA levels to promote meiosis.” Nat Commun 10(1): 3414.

Carpenter, K., R. B. Bell, J. Yunus, A. Amon and L. E. Berchowitz (2018). “Phosphorylation-Mediated Clearance of Amyloid-like Assemblies in Meiosis.” Dev Cell 45(3): 392–405 e396.

Chen, X., R. Gaglione, T. Leong, L. Bednor, T. de Los Santos, E. Luk, M. Airola and N. M. Hollingsworth (2018). “Mek1 coordinates meiotic progression with DNA break repair by directly phosphorylating and inhibiting the yeast pachytene exit regulator Ndt80.” PLoS Genet 14(11): e1007832.

Clancy, M. J., M. E. Shambaugh, C. S. Timpte and J. A. Bokar (2002). “Induction of sporulation in Saccharomyces cerevisiae leads to the formation of N6-methyladenosine in mRNA: a potential mechanism for the activity of the IME4 gene.” Nucleic acids research 30(20): 4509–4518.

Clifford, D. M., S. M. Marinco and G. S. Brush (2004). “The meiosis-specific protein kinase Ime2 directs phosphorylation of replication protein A.” J Biol Chem 279(7): 6163–6170.

Collier, B., B. Gorgoni, C. Loveridge, H. J. Cooke and N. K. Gray (2005). “The DAZL family proteins are PABP-binding proteins that regulate translation in germ cells.” EMBO J 24(14): 2656–2666.

Corbi, D., S. Sunder, M. Weinreich, A. Skokotas, E. S. Johnson and E. Winter (2014). “Multisite phosphorylation of the Sum1 transcriptional repressor by S-phase kinases controls exit from meiotic prophase in yeast.” Mol Cell Biol 34(12): 2249–2263.

Deng, C. and W. S. Saunders (2001). “RIM4 encodes a meiotic activator required for early events of meiosis in Saccharomyces cerevisiae.” Mol Genet Genomics 266(3): 497–504.

Dirick, L., L. Goetsch, G. Ammerer and B. Byers (1998). “Regulation of meiotic S phase by Ime2 and a Clb5,6-associated kinase in Saccharomyces cerevisiae.” Science 281(5384): 1854–1857.

Holt, L. J., J. E. Hutti, L. C. Cantley and D. O. Morgan (2007). “Evolution of Ime2 phosphorylation sites on Cdk1 substrates provides a mechanism to limit the effects of the phosphatase Cdc14 in meiosis.” Mol Cell 25(5): 689–702.

Jin, L., K. Zhang, Y. Xu, R. Sternglanz and A. M. Neiman (2015). “Sequestration of mRNAs Modulates the Timing of Translation during Meiosis in Budding Yeast.” Mol Cell Biol 35(20): 3448–3458.

Kassir, Y., D. Granot and G. Simchen (1988). “IME1, a positive regulator gene of meiosis in S. cerevisiae.” Cell 52(6): 853–862.

Kurihara, L. J., C. T. Beh, M. Latterich, R. Schekman and M. D. Rose (1994). “Nuclear congression and membrane fusion: two distinct events in the yeast karyogamy pathway.” The Journal of cell biology 126(4): 911–923.

Kurihara, L. J., B. G. Stewart, A. E. Gammie and M. D. Rose (1996). “Kar4p, a karyogamy-specific component of the yeast pheromone response pathway.” Mol Cell Biol 16(8): 3990–4002.

Lahav, R., A. Gammie, S. Tavazoie and M. D. Rose (2007). “Role of transcription factor Kar4 in regulating downstream events in the Saccharomyces cerevisiae pheromone response pathway.” Mol Cell Biol 27(3): 818–829.

Laureau, R., A. Dyatel, G. Dursuk, S. Brown, H. Adeoye, J. X. Yue, M. De Chiara, A. Harris, E. Unal, G. Liti, I. R. Adams and L. E. Berchowitz (2021). “Meiotic Cells Counteract Programmed Retrotransposon Activation via RNA-Binding Translational Repressor Assemblies.” Dev Cell 56(1): 22–35 e27.

Lin, S., J. Choe, P. Du, R. Triboulet and R. I. Gregory (2016). “The m(6)A Methyltransferase METTL3 Promotes Translation in Human Cancer Cells.” Mol Cell 62(3): 335–345.

McIsaac, R. S., P. A. Gibney, S. S. Chandran, K. R. Benjamin and D. Botstein (2014). “Synthetic biology tools for programming gene expression without nutritional perturbations in Saccharomyces cerevisiae.” Nucleic Acids Res 42(6): e48.

Monteiro, P. T., J. Oliveira, P. Pais, M. Antunes, M. Palma, M. Cavalheiro, M. Galocha, C. P. Godinho, L. C. Martins, N. Bourbon, M. N. Mota, R. A. Ribeiro, R. Viana, I. Sa-Correia and M. C. Teixeira (2020). “YEASTRACT+: a portal for cross-species comparative genomics of transcription regulation in yeasts.” Nucleic Acids Res 48(D1): D642–D649.

Neiman, A. M. (2011). “Sporulation in the budding yeast Saccharomyces cerevisiae.” Genetics 189(3): 737–765.

Pak, J. and J. Segall (2002). “Regulation of the premiddle and middle phases of expression of the NDT80 gene during sporulation of Saccharomyces cerevisiae.” Mol Cell Biol 22(18): 6417–6429.

Schwartz, S., S. D. Agarwala, M. R. Mumbach, M. Jovanovic, P. Mertins, A. Shishkin, Y. Tabach, T. S. Mikkelsen, R. Satija, G. Ruvkun, S. A. Carr, E. S. Lander, G. R. Fink and A. Regev (2013). “High-resolution mapping reveals a conserved, widespread, dynamic mRNA methylation program in yeast meiosis.” Cell 155(6): 1409–1421.

Sedgwick, C., M. Rawluk, J. Decesare, S. Raithatha, J. Wohlschlegel, P. Semchuk, M. Ellison, J. Yates, 3rd and D. Stuart (2006). “Saccharomyces cerevisiae Ime2 phosphorylates Sic1 at multiple PXS/T sites but is insufficient to trigger Sic1 degradation.” Biochem J 399(1): 151–160.

Shin, M. E., A. Skokotas and E. Winter (2010). “The Cdk1 and Ime2 protein kinases trigger exit from meiotic prophase in Saccharomyces cerevisiae by inhibiting the Sum1 transcriptional repressor.” Mol Cell Biol 30(12): 2996–3003.

Soushko, M. and A. P. Mitchell (2000). “An RNA-binding protein homologue that promotes sporulation-specific gene expression inSaccharomyces cerevisiae.” Yeast 16(7): 631–639.

van Werven, F. J. and A. Amon (2011). “Regulation of entry into gametogenesis.” Philos Trans R Soc Lond B Biol Sci 366(1584): 3521–3531.

Wang, F., R. Zhang, W. Feng, D. Tsuchiya, O. Ballew, J. Li, V. Denic and S. Lacefield (2020). “Autophagy of an Amyloid-like Translational Repressor Regulates Meiotic Exit.” Dev Cell 52(2): 141–151 e145.

Wang, P., K. A. Doxtader and Y. Nam (2016). “Structural Basis for Cooperative Function of Mettl3 and Mettl14 Methyltransferases.” Mol Cell 63(2): 306–317.

Wang, X., J. Feng, Y. Xue, Z. Guan, D. Zhang, Z. Liu, Z. Gong, Q. Wang, J. Huang, C. Tang, T. Zou and P. Yin (2016). “Structural basis of N(6)-adenosine methylation by the METTL3-METTL14 complex.” Nature 534(7608): 575–578.

Wei, X., Y. Huo, J. Pi, Y. Gao, S. Rao, M. He, Q. Wei, P. Song, Y. Chen, D. Lu, W. Song, J. Liang, L. Xu, H. Wang, G. Hong, Y. Guo, Y. Si, J. Xu, X. Wang, Y. Ma, S. Yu, D. Zou, J. Jin, F. Wang and J. Yu (2022). “METTL3 preferentially enhances non-m(6)A translation of epigenetic factors and promotes tumourigenesis.” Nat Cell Biol 24(8): 1278–1290.

Winter, E. (2012). “The Sum1/Ndt80 transcriptional switch and commitment to meiosis in Saccharomyces cerevisiae.” Microbiol Mol Biol Rev 76(1): 1–15.

